# The conserved two-component systems CutRS and CssRS control the protein secretion stress response in Streptomyces species

**DOI:** 10.1101/2025.03.20.644377

**Authors:** Thomas C. McLean, Ainsley Beaton, Neil A. Holmes, Carlo Martins, Gerhard Saalbach, Govind Chandra, Sibyl F.D. Batey, Jana China, Barrie Wilkinson, Matthew I. Hutchings

**Author notes:** These authors contributed equally to this work (Thomas C. McLean, Ainsley Beaton).

## Abstract

*Streptomyces* secondary metabolites account for over half of all clinically used antibiotics, as well as numerous antifungal agents, anticancer compounds, and immunosuppressants. Two-component systems, which are widespread in bacteria, are key regulators of antibiotic production in *Streptomyces* species, yet their activating signals remain poorly understood. CutRS was the first two-component system identified in the genus *Streptomyces* and deletion of *cutRS* in *Streptomyces coelicolor* was shown to enhance antibiotic production, although its CutR regulon does not include any biosynthetic genes. Here, we used *Streptomyces venezuelae* to further investigate CutRS function. We show that deletion of *cutRS* leads to an increase in growth rate and a reversal of the typical glucose-mediated carbon catabolite repression typically observed in *Streptomyces* species. We also demonstrate that CutR DNA binding is glucose-dependent, but CutR does not directly regulate genes involved in growth, antibiotic biosynthesis, or glucose metabolism. The only CutR targets conserved in both *S. coelicolor* and *S. venezuelae* are the foldase genes *htrA3* and *htrB*, which are involved in the protein secretion stress response. Consistent with this, we show that CutS homologues all contain two conserved cysteine residues in their extracellular sensor domains and that changing these residues to serine constitutively activates *S. venezuelae* CutRS. We propose that failure of a disulfide bond to form between these cysteine residues indicates secretion stress and leads to activation of the CutRS system and the secretion stress response.

**IMPORTANCE:** *Streptomyces* bacteria are the primary source of clinically useful antibiotics. While many two-component systems have been linked to antibiotic biosynthesis in *Streptomyces* species, few have been well characterized. Here, we characterize a secretion stress sensing two-component system called CutRS and propose a model for how the sensor kinase detects extracellular protein misfolding via two highly conserved cysteine residues. Importantly, we also show that deletion of *cutRS* triggers antibiotic overproduction in the presence of glucose. Since glucose normally represses antibiotic biosynthesis in *Streptomyces* species through carbon catabolite repression, this finding reveals a simple genetic route to bypass this barrier. This has significant implications for antibiotic discovery pipelines and industrial production, where glucose-rich media are preferred for cost and scalability. Our results position CutRS as a key target for future strain improvement strategies.

## INTRODUCTION

*Streptomyces* species are studied for their complex life cycles and prolific production of specialized metabolites, including many clinically important antibiotics (1). These metabolites are typically encoded by biosynthetic gene clusters (BGCs), but despite the wealth of genomic data available, only ∼3% of BGCs have been linked to their corresponding natural products (2). The vast majority remain cryptic under standard laboratory conditions, representing untapped reservoirs for novel bioactive compounds (3, 4). Unlocking these cryptic BGCs requires a better understanding of the regulatory mechanisms, including signal transduction pathways, that control specialized metabolism.

BGC expression is often regulated by transcription factors encoded within the cluster itself (cluster-situated regulators), as well as by global regulatory systems that integrate signals related to nutrient availability, growth phase, and environmental stress to specialized metabolite production (5). Among these global regulators, two-component systems (TCSs) are particularly important. TCSs are widespread in bacteria, fungi, and plants, and many have been shown to influence antibiotic production in *Streptomyces* species. However, the mechanisms by which they exert this control are often poorly understood (6–9).

TCSs typically consist of a membrane-bound sensor kinase (SK) and a cognate response regulator (RR). Upon detection of an external signal, the SK autophosphorylates and transfers the phosphate to the RR, which then dimerizes and binds DNA to modulate the transcription of target genes. *Streptomyces* genomes encode between 50 and 100 TCSs, yet only 15 are highly conserved across species (10). Of these, several have been shown to impact antibiotic biosynthesis, but only a small number have been functionally characterized, including their target regulons.

CutRS was the first TCS identified in the genus *Streptomyces* (11), and its disruption in *S. lividans* and *S. coelicolor* resulted in overproduction of the antibiotic actinorhodin (11, 12). However, a subsequent study in *S. coelicolor* revealed that none of the 88 genes directly regulated by CutR were involved in actinorhodin biosynthesis (13). Instead, the CutRS system was implicated in the protein secretion stress response, with CutR activating *htrA3* and repressing *htrB*, both of which encode high-temperature requirement (Htr) chaperone-proteases that assist in folding or degradation of misfolded proteins in the extracellular space (13).

Here, we investigated the CutRS system in *Streptomyces venezuelae*, a genetically tractable and rapidly growing model organism that has emerged as a powerful system for studying *Streptomyces* development and regulation (14). We found that *S. venezuelae* CutR binds nine promoter regions, but the core CutR regulon shared with *S. coelicolor* includes only *htrA3* and *htrB*, reinforcing a conserved role in secretion stress regulation. Alignment of >100 CutS homologues across *Streptomyces* species revealed two invariant cysteine residues as the only conserved features in their predicted extracellular sensor domains. We demonstrate that changing these cysteines to serine constitutively activates CutRS, suggesting that the system senses protein misfolding via disulfide bond status outside the cell. Although CutRS is only conserved in *Streptomyces* species, analysis of SK sequences from 12,799 bacterial genomes revealed that 99% encode at least one SK with two or more cysteines in their predicted extracellular sensor domains. We therefore propose that extracellular redox sensing could be a widespread signaling mechanism across the bacterial domain.

## RESULTS

### Deletion of cutRS affects growth and antibiotic production in S. venezuelae

To investigate the function of CutRS, we deleted the *cutRS* operon in *S. venezuelae* and found that the Δ*cutRS* mutant exhibits a striking phenotype when grown on yeast peptone + D-glucose (YPD) agar. The mutant displays increased antibacterial activity and accelerated growth (Figures 1, 2A, and 2B). Further analysis revealed that the Δ*cutRS* strain produces 10-fold more biomass than the wild-type strain on YPD agar. This phenotype was not restricted to YPD medium; a similar increase in biomass was observed on MYM agar, but again only in the presence of D-glucose (Figure 2B). This increased growth correlates with the rapid depletion of D-glucose from the medium (Figure 2C). Consistent with this, in the absence of D-glucose (YP agar), no phenotypic differences were observed between the wild-type and Δ*cutRS* strains (Figures 1A, 2A, and 2B).

**Figure 1.**
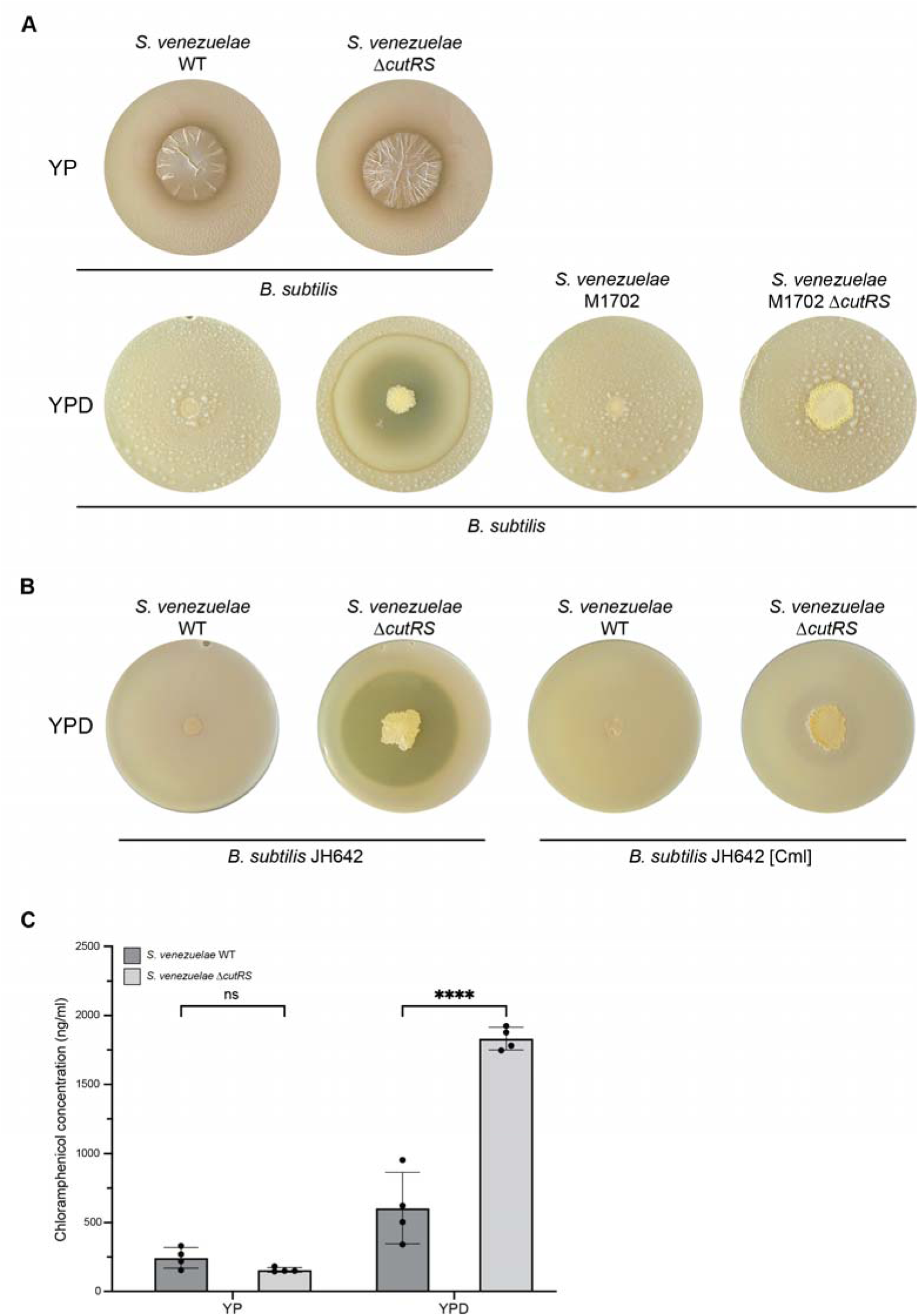
| Deletion of cutRS results in the overproduction of chloramphenicol. A. Antimicrobial overlay assays of the S. venezuelae wild-type, ΔcutRS, M1702 (ΔcmlΔjad) and M1702 ΔcutRS strains with B. subtilis. B. Antimicrobial overlay assays of S. venezuelae wild-type and ΔcutRS on YPD agar. (left) Overlays with the chloramphenicol-sensitive strain B. subtilis JH642. (right) Overlays with the chloramphenicol-resistant strain B. subtilis JH642 Cml^R^ C. Quantification of chloramphenicol extracted from S. venezuelae wild-type and ΔcutRS strains on YP and YPD agar by HPLC.

**Figure 2.**
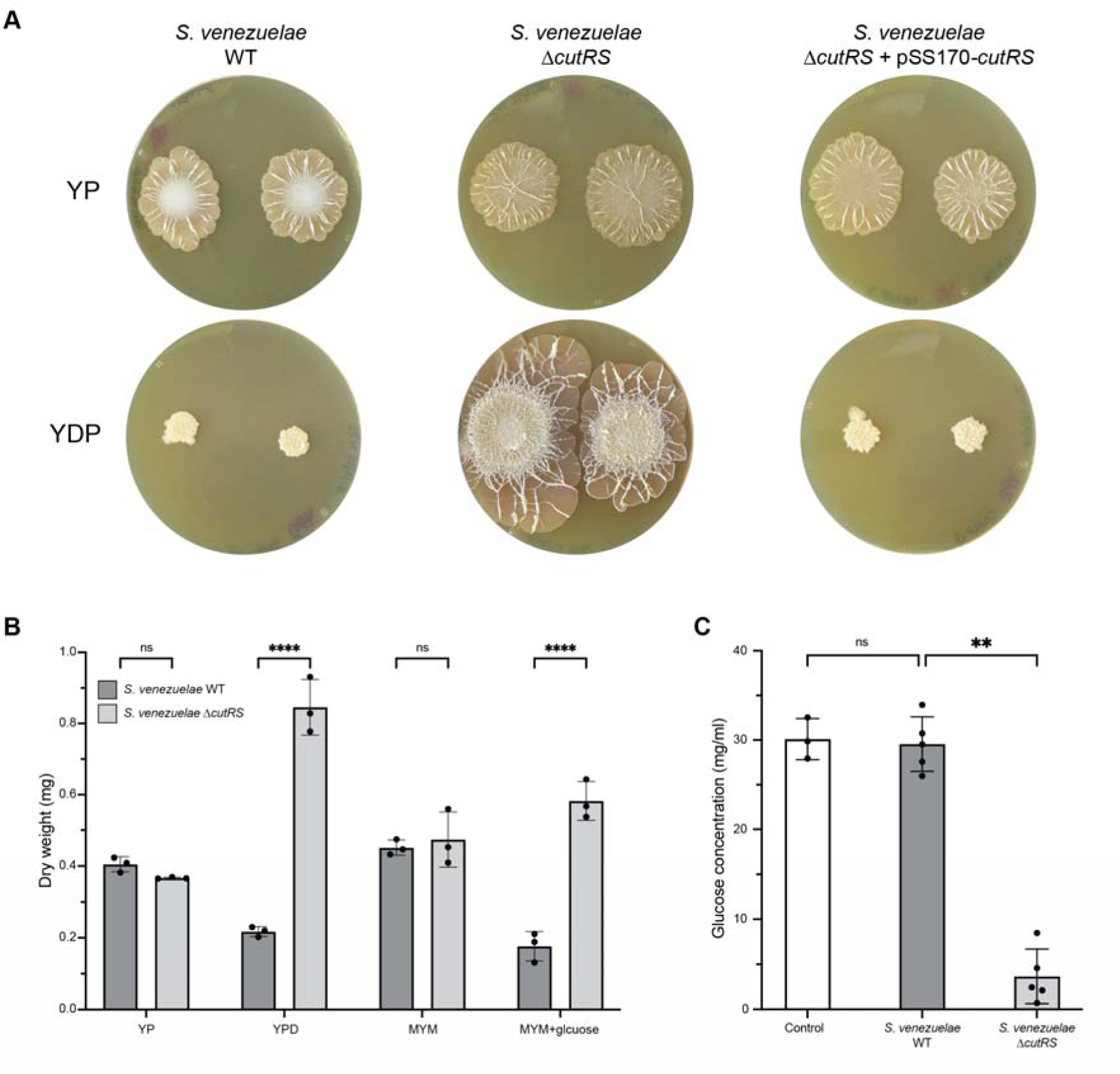
| The CutRS two-component system is involved in colony growth and glucose consumption. A. S. venezuelae wild-type, ΔcutRS and ΔcutRS +pSS170-cutRS colonies grown on YP (-glucose) and YPD (+ glucose) agar for 10 days at 30°C. B. The colony dry weight (mg) of S. venezuelae wild-type and ΔcutRS after 10 days of growth at 30°C on YP, YPD, MYM and MYM+glucose. C. The remaining glucose concentration (mg/ml) in spent media after 10 days of growth at 30°C (S. venezuelae wild-type or ΔcutRS).

When challenged with *Bacillus subtilis* on YP agar, both the wild-type and Δ*cutRS* strains produced small zones of inhibition (Figure 1A). However, on YPD agar, bioactivity was lost in the wild-type strain, consistent with the known inhibition of antibiotic biosynthesis by D-glucose via carbon catabolite repression (15), while the Δ*cutRS* strain exhibited a markedly larger zone of inhibition (Figure 1A). This suggests that D-glucose triggers antibiotic overproduction in the Δ*cutRS* strain, supporting previous observations that glucose activates actinorhodin production in the *S. coelicolor* Δ*cutRS* mutant (13). Notably, this is atypical, as D-glucose generally represses antibiotic production in *Streptomyces* species (15, 16).

*S. venezuelae* is known to produce chloramphenicol and jadomycin, both of which inhibit *B. subtilis*. To determine which antibiotic was responsible for the observed activity, we deleted both BGCs to generate strain M1702. This strain failed to inhibit *B. subtilis* growth, indicating that the bioactivity of the parent strain was due to either chloramphenicol or jadomycin (Figure 1A). To further resolve this, we challenged the wild-type and Δ*cutRS* strains with a chloramphenicol-resistant *B. subtilis* strain. The Δ*cutRS* strain produced only a faint zone of inhibition, markedly smaller than that observed against the chloramphenicol-sensitive strain, confirming that the bioactivity was primarily due to chloramphenicol (Figure 1B).

Finally, we extracted cultures from YPD agar plates with ethyl acetate and analyzed the 120 samples by HPLC. The Δ*cutRS* strain produced approximately three times more chloramphenicol than the wild-type in the presence of D-glucose, confirming that increased chloramphenicol production is responsible for the enhanced antibacterial activity of the Δ*cutRS* strain (Figure 1C).

### Identifying CutR target promoters and binding sites in S. venezuelae

Two-component systems typically regulate gene expression through their response regulators (RRs)—in this case, CutR. To define the CutR regulon, we complemented the *S. venezuelae* Δ*cutRS* strain with a modified *cutRS* operon encoding wild-type CutS and a C-terminally 3×Flag-tagged CutR. We then cultured both the wild-type and Flag-tagged strains on YP and YPD agar and performed ChIP-seq using monoclonal α-Flag antibodies.

Analysis of the sequencing data (accession number GSE225370) revealed a single significantly enriched peak in the Flag-tagged CutR strain grown on YP agar (absence of D-glucose), located upstream of the *cutRS* operon. In contrast, ten enriched peaks were detected in cultures grown on YPD agar (presence of D-glucose) (Table 1). These included peaks upstream of *htrA3* and *htrB*, which are also known CutR targets in *S. coelicolor* (Figure 3A), as well as a strong peak upstream of *vnz_08815*, a putative cell wall amidase gene.

**Figure 3.**
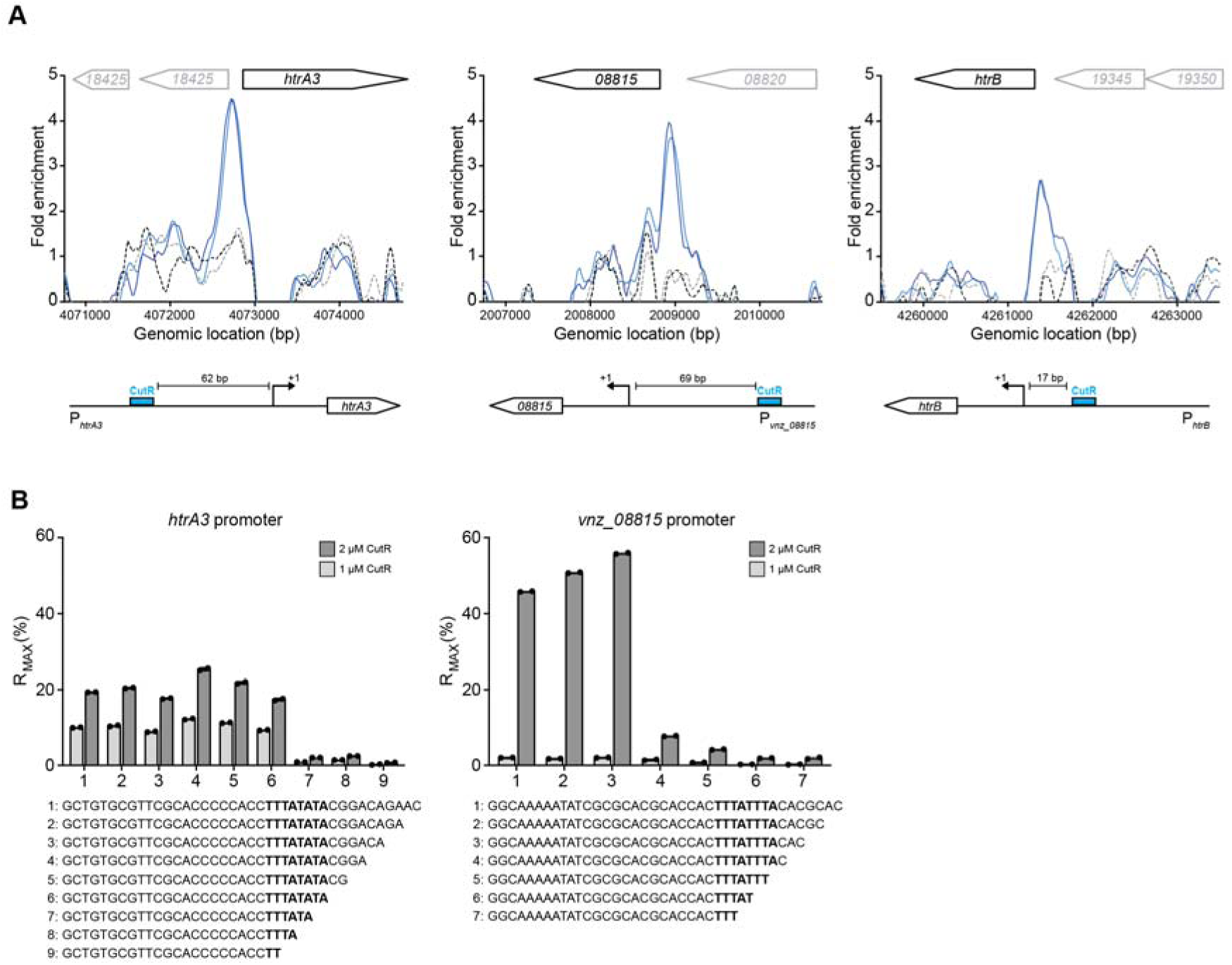
| The defined DNA binding site for the CutR response regulator. A. ChIP-seq peaks for biological replicates of CutR-3xFLAG (blue lines) and the wild-type control (grey lines). CutR binds upstream of the genes htrA3, vnz_08815 and htrB. B. ReDCaT SPR was used to determine the binding site of 6xHis-CutR to the promoter regions of htrA3 and the putative target gene vnz_08815. Sequential truncations of 2 bp were used to define the precise binding motif of TAWATAAA. The %R_MAX_ was determined using the molecular masses of the annealed oligo duplex and monomeric 6xHis-CutR. This binding motif was then used to identify the CutR binding site upstream of htrB (A).

**Table 1.** Gene promoters with CutR binding sites that are enriched in the ChIP-seq dataset. The binding site positions are given relative to the transcript start sites (TSS). The DNA sequences of the CutR binding sites are shown, two of which (htrA3 and vnz_08815) were verified in vitro using ReDCaT SPR (Figure 3).

| Gene | Predicted function | CutR binding site relative to TSS | Binding site sequence | Predicted effect | Protein abundance (WT/$\Delta cutRS$) |
| --- | --- | --- | --- | --- | --- |
| *vnz_06385* | Acyl coA dehydrogenase | +21 | TAATTAAA | Repression | 0.795 |
| *vnz_06390* | TetR-family regulator | +28 | TAATTAAA | Repression | 0.947 |
| *vnz_08815* | Cell wall amidase | -69 | TAAATAAA | Activation | 2.56 |
| *vnz_12615* | Membrane protein | -4 | TACATAAA | Repression | ND |
| *htrA3* | Quality control protease | -62 | TATATAAA | Activation | 8.134 |
| *htrB* | Quality control protease | -17 | CATATAAA | Repression | 0.053 |
| *vnz_29845* | NAD(P)/FAD-dependent oxidoreductase | -28 | CAAATAAA | Repression | 0.707 |
| *vnz_33990* | Beta mannosidase | -21 | TAAGTAAA | Repression | 0.747 |
| *vnz_33995* | Sugar binding protein for ABC transport | +1 | TAAGTAAA | Repression | 0.977 |

To precisely define CutR binding sites, we expressed and purified hexahistidine-tagged CutR for use in a DNA footprinting method called Reusable DNA Capture Technology Surface Plasmon Resonance (ReDCaT SPR) (17). Using tiled, double-stranded oligonucleotide probes spanning the promoters of *htrA3* and *vnz_08815*, we measured CutR binding affinity via SPR. A preliminary scan identified the core binding regions, and these probes were then truncated in 2 bp steps to determine the minimal binding sequences (Figure 3B). CutR bound the sequence TATATAAA in the *htrA3* promoter and TAAATAAA in the *vnz_08815* promoter (Figure 3B). From these and other enriched regions, we defined the CutR consensus binding sequence as TAWATAAA.

### CutR is both a transcriptional activator and repressor

Having identified CutR binding sites at its target promoters (Table 1), we cross-referenced them with publicly available *S. venezuelae* transcription start site (TSS) data from http://streptomyces.org.uk (18). The CutR binding site at the *htrB* promoter is located 17 bp upstream of the TSS, while at the *htrA3* promoter it lies 62 bp upstream (Table 1 and Figure 3A). The position of a regulator binding site relative to the TSS often determines whether it functions as an activator or repressor (19). Binding 17 bp upstream of the *htrB* TSS likely overlaps the core promoter region, interfering with RNA polymerase binding and resulting in transcriptional repression (20, 21). In contrast, binding further upstream, such as at the *htrA3* promoter, may facilitate transcriptional activation via interaction between CutR and RNA polymerase bound at the –35 site (21–23).

To assess the effect of *cutRS* deletion on CutR target gene products, we used tandem mass tag (TMT) quantitative proteomics. Of the ten predicted CutR-dependent proteins, six were detected (Table 1). These data show that CutR activates expression of HtrA3 (8-fold increase in the wild-type strain compared to the *ΔcutRS* mutant) and the putative cell wall amidase Vnz_08815 (2.5-fold increase), consistent with the location of its binding sites upstream of their promoters (Figure 4, Table 1, and Supplementary Table 1). In contrast, HtrB production was repressed by CutR, showing a 19-fold increase in the Δ*cutRS* strain relative to the wild-type, consistent with CutR binding in a position that occludes RNA polymerase binding (Figure 3A).

**Figure 4.**
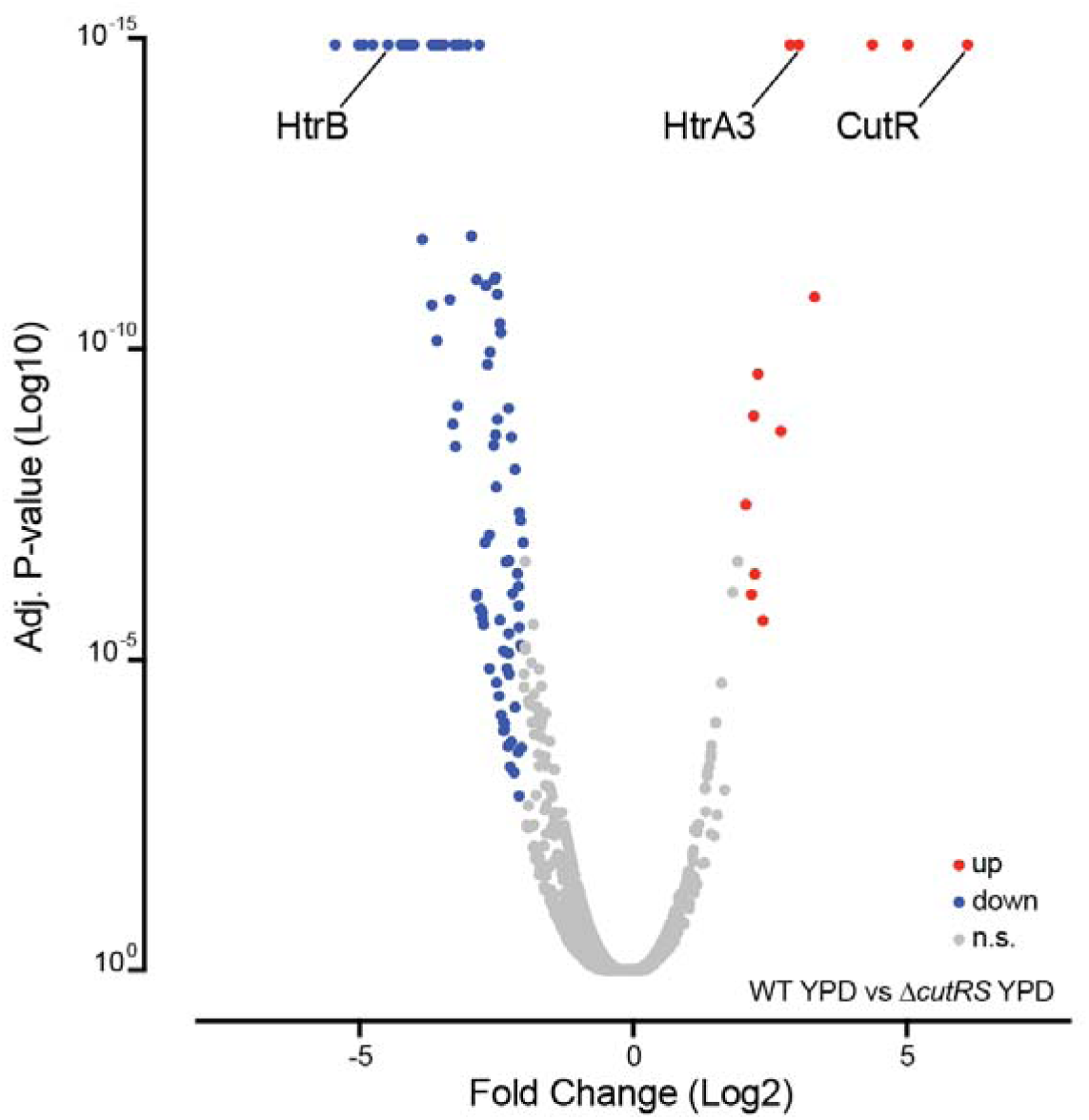
| The deletion of cutRS causes a global shift in the proteome. Volcano plot of the Fold Change (Log_2_) protein abundance detected by TMT-proteomics against the calculated adjusted P-value (Log_10_) for S. venezuelae wild-type (wild-type) and ΔcutRS strains on YPD agar. The abundance of CutRS in the wild-type samples appears exaggerated due to the data processing against the ΔcutRS samples and does not represent normal levels.

The expression of the remaining detected target genes was not significantly affected by *cutRS* deletion. This may be because these genes are not directly regulated by CutR under the tested growth conditions, because CutR binding does not alter transcription in all cases, or because additional layers of regulation are involved (Table 1).

### CutS activity is controlled by two conserved cysteines in its extracellular sensor domain

The conservation of *htrA3* and *htrB* as CutR targets in the distantly related species *S. coelicolor* and *S. venezuelae* supports the hypothesis that CutRS plays a role in the protein secretion stress response. This further implies that CutRS complements the function of CssRS, which activates the expression of *htrA1*, *htrA2* and *htrB*, but not *htrA3* (24). We thus set out to investigate how the extracellular sensor domain, which is predicted to be between the two transmembrane helices of CutS, might detect defects in protein folding on the outside of the cell. We first aligned the extracellular sensor domains of >100 CutS proteins taken from complete, published *Streptomyces* genome sequences to look for conserved residues or motifs. This analysis identified two cysteine residues that are conserved across every CutS sensor domain (Supplementary Figure 3). These are the only conserved residues in the CutS sensor domain. Cysteines are important in extracellular protein folding in actinobacteria (25) and, consistent with this, the analysis of all proteins encoded in *S. venezuelae* with a Sec signal sequence revealed that 74% contain two or more cysteines that likely form disulfide bonds to help them fold. We thus hypothesized that CutS monitors disulfide bond formation in Sec-translocated proteins. Given the conservation of the two cysteine residues, we reasoned that this could be via the making or breaking of a disulfide bond in the CutS extracellular sensor domain.

Modelling of the *S. venezuelae* CutS extracellular sensor domain using AlphaFold indicated that these cysteine residues likely sit within 5Å of each other, the distance required to form disulfide bonds (Supplementary Figure 4). To test this, we designed a CutS variant where both cysteine residues were replaced with serine (CutS(C85S,C103S)) and introduced it into the *S. venezuelae* wild-type and Δ*cutRS* strains along with the wild-type *cutR* gene. This restored a wild-type growth phenotype to the Δ*cutRS* mutant (Figure 5A) which suggests the CutS(C85S,C103S) variant is active. However, previous work on the VanRS TCS demonstrated that it is possible for loss of a SK to render its cognate RR constitutively active (26). To discount this possibility, we introduced individual copies of either *cutS* or *cutR* under the control of the *ermE*\* promoter into the Δ*cutRS* mutant, but neither restored the wild-type phenotype, confirming that CutS is required for CutR activity. To further investigate the activity of the CutS(C85S,C103S) variant we used qRT-PCR to measure the expression of the CutR target genes *htrA3* and *htrB* in the wild-type, Δ*cutRS* and CutS(C85S,C103S) strains (Figure 5B). The expression of *htrA3* was higher in the strain harboring the CutS(C85S,C103S) variant compared to the wild-type strain, while the expression of *htrB* was reduced (Figure 5B) which is consistent with CutS(C85S,C103S) activating CutR which then directly activates the transcription of *htrA3* and represses *htrB*. Thus, we conclude that replacing the two extracellular sensor domain cysteines with serine residues results in a constitutively active CutRS system which supports the hypothesis that an inability to form disulfide bonds outside the cell activates CutRS and triggers the secretion stress response.

**Figure 5.**
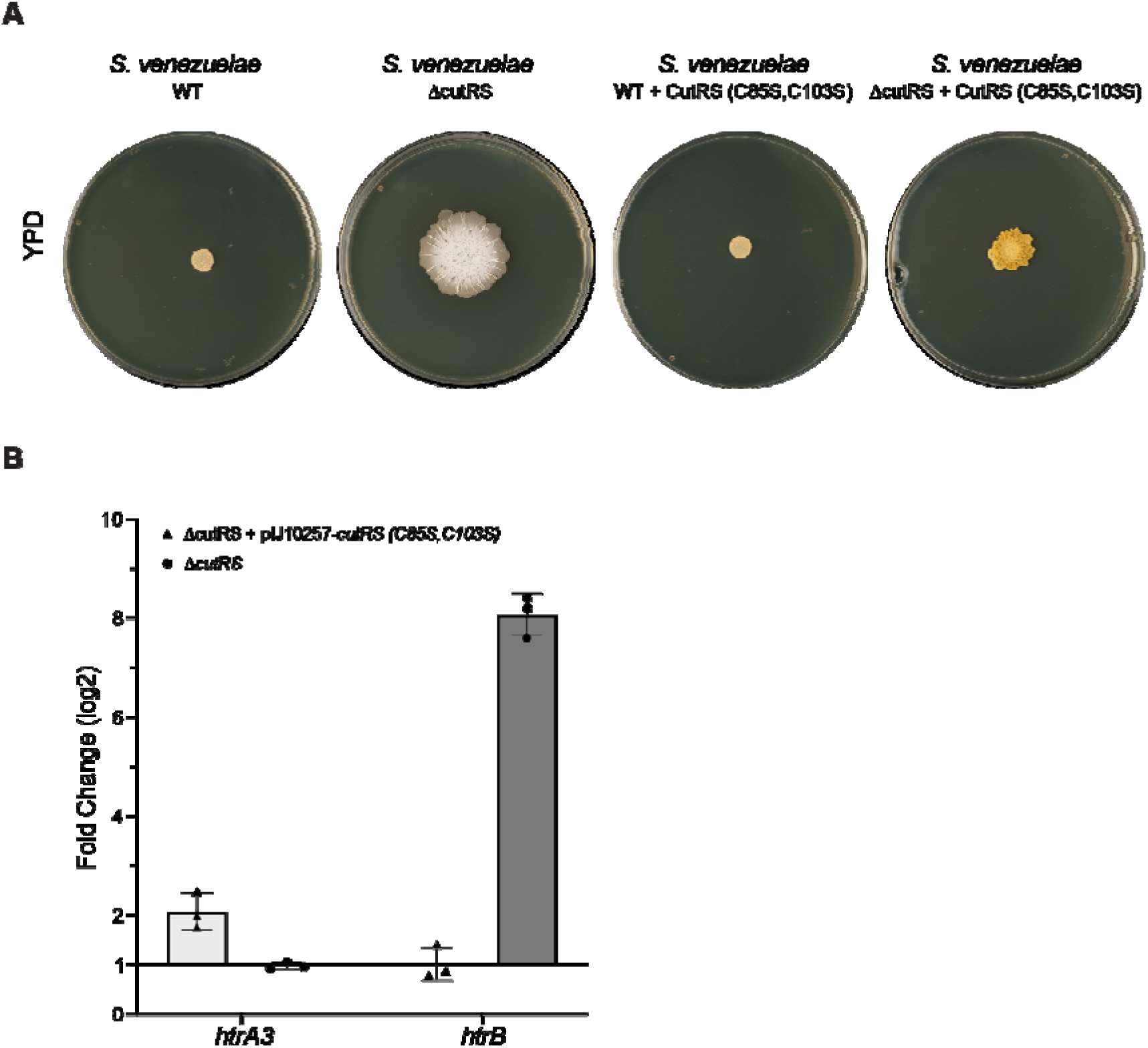
| The conserved extracellular cysteine residues control the activity of CutS. A. The ΔcutRS mutant can be partially complemented by in trans expression of an operon encoding wild-type CutR and a CutS protein in which the extracellular cysteines are altered to serine (CutS(C85S,C103S)). B. Quantitative RT-PCR analyzing the abundance of htrA3 (light grey bars) and htrB (dark grey bars) transcripts normalized to constitutively expressed hrdB and compared to wild-type. Error bars represent standard deviation across replicates.

### The wild-type phenotype is recovered by the deletion of cssRS in the ΔcutRS background

This work and previous reports suggest that the secretion stress response in *Streptomyces* species is controlled by the CssRS and CutRS two-component systems which together control the expression of *htrA1*, *htrA2*, *htrA3* and *htrB*, which encode the four conserved HtrA-family foldases in *Streptomyces* species (24, 27). Although the CssR regulon has not yet been defined, electrophoretic mobility shift assays alongside qRT-PCR were used previously in *S. lividans* (a very close relative of *S. coelicolor*) to show that CssR directly activates expression of *htrA1*, *htrA2* and *htrB*, but not *htrA3* (24). Instead, the expression of *htrA3* is directly activated by CutR, which also represses the expression of *htrB* in *S. coelicolor* and *S. venezuelae* (27). We attempted to replicate the Δ*cutRS* phenotype by deleting *htrA3* and overexpressing *htrB* in the same background (Supplementary Figure 2), but this did not result in the phenotypic change on YPD that we observed with Δ*cutRS*. We thus reasoned that the CssRS system might be compensating for the loss of CutRS. To further investigate the interplay of CssRS and CutRS we made single and double Δ*cssRS* and Δ*cutRS* mutants of *S. venezuelae* and observed their phenotypes on YP and YPD agar. In the absence of D-glucose, the wild-type, Δ*cssRS*, Δ*cutRS* and Δ*cssRS-*Δ*cutRS* strains all displayed the same phenotype (Figure 6). On YPD agar, growth is reduced in the wild-type strain by the addition of D-glucose to the medium, but the Δ*cutRS* mutant consumes the glucose and reverts to a phenotype like that observed on YP agar (Figures 1 and 5). Deletion of *cssRS* in the wild-type background has no obvious effect on growth on either YP or YPD agar, but deletion of *cssRS* in the Δ*cutRS* mutant restores the wild-type phenotype to the *ΔcutRS* mutant growing on YPD agar, suggesting that the altered growth of the Δ*cutRS* mutant on YPD is dependent on CssRS (Figure 6). Proteomics analysis of the wild-type and Δ*cutRS* strains revealed that the levels of CssRS are increased 5-fold in the Δ*cutRS* mutant, and as previously discussed, HtrB levels (activated by CssR and repressed by CutR) are increased 20-fold in the absence of *cutRS*. The *htrB* and *cssRS* genes are located adjacent to each other and likely form an operon under the direct control of both CutR (negative) and CssR (positive). Overproduction of HtrB could be due to the loss of CutR repression in the mutant strain combined with the overproduction of CssRS, with CssR further activating *htrB* expression. A future challenge will be to define the CssR regulon and determine how deletion of *cssRS* complements Δ*cutRS*.

**Figure 6.**
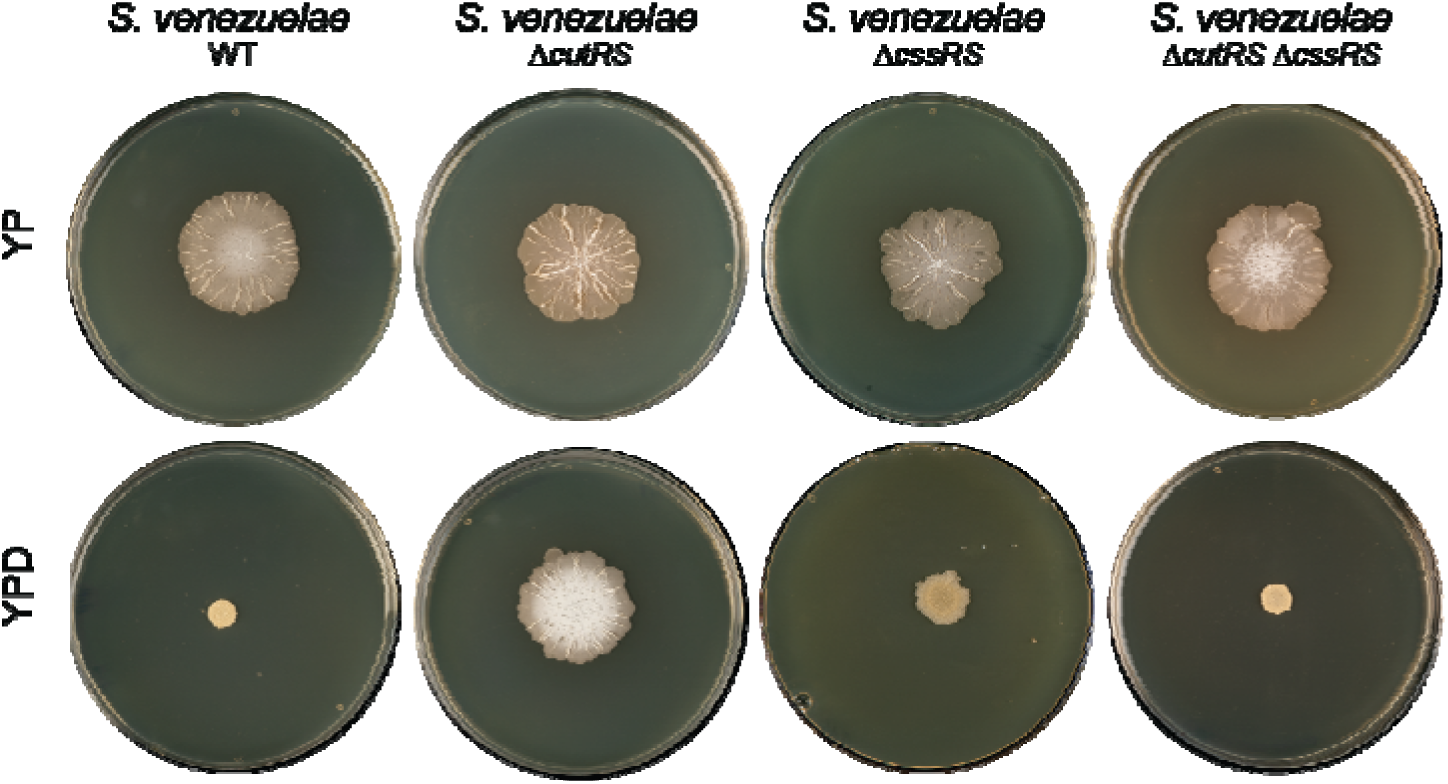
| The S. venezuelae wild-type phenotype is recovered by deletion of cssRS in the ΔcutRS background. S. venezuelae wild-type, ΔcutRS, ΔcssRS and ΔcutRS ΔcssRS strains grown on YP (-glucose) and YPD (+ glucose) agar for 10 days at 30°C.

### Extracellular disulfide sensing could be conserved across bacteria

Analysis of the 15 conserved two-component systems in *Streptomyces* species (28–30) revealed that only CutS and CssS have two conserved extracellular cysteines. Given that these cysteines are implicated in the control of CutS activity we hypothesise the same may be true for CssS and perhaps other bacterial SKs. To investigate how widespread this might be in the bacterial kingdom, we analyzed 12,799 high-quality bacterial genomes, identifying 348,633 proteins predicted to function as SKs. Using DeepTMHMM (31) we predicted the transmembrane helices for all the SKs and then used this information to identify the putative sensor domains of these proteins and quantify the number of cysteine residues present in these extracellular domains. We observed that 98.9% of all the bacterial strains examined encode at least one SK with two or more cysteines in their predicted extracellular sensor domains (Figure 7A).

**Figure 7.**
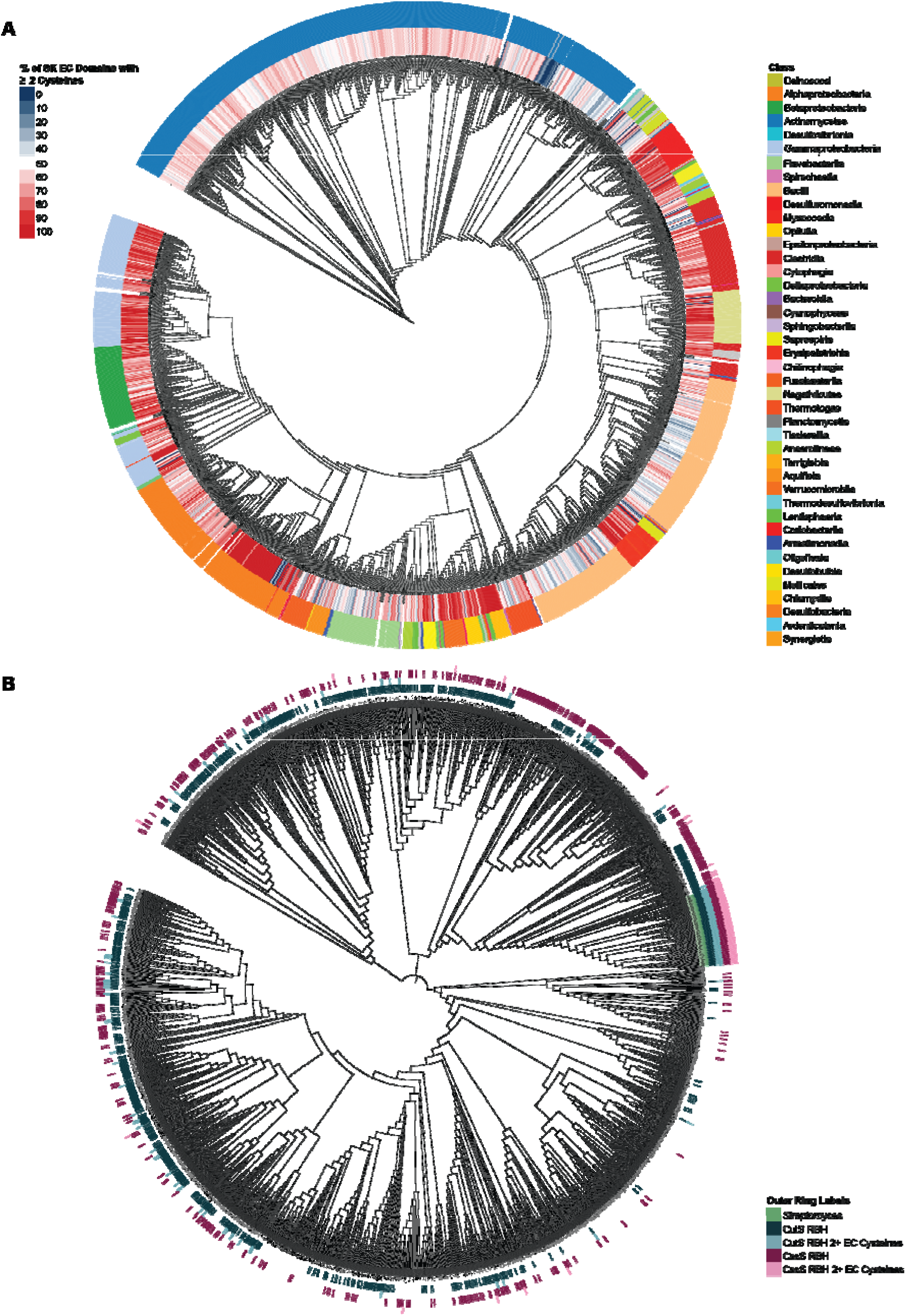
| Phylogenetic and cysteine residue analysis of bacterial sensor kinases. A. Maximum likelihood phylogenetic tree of 348,633 predicted SKs from 12,799 high-quality bacterial genomes. The outer ring is color-coded by bacterial class, as indicated in the legend. The heatmap overlay in the inner ring shows the proportion of SKs with two or more cysteine residues in their extracellular domains, as predicted by DeepTMHMM. Dark blue indicates 0% of the SKs in that genome can form a disulphide bond in their extracellular domain, increasing to dark red, which indicates 100% of the SKs in that genome can form a disulphide bond in their extracellular domain. B. Maximum likelihood phylogenetic tree based on RpoB protein sequences from 2,940 bacterial reference genomes. The outer rings represent the following: Genomes in the Streptomyces genus (light green); CutS reciprocal BLAST best hits (RBBH) (dark blue); CutS homologs with two or more cysteine residues in their extracellular domains, as predicted by DeepTMHMM (light blue); CssS reciprocal BLAST best hits (RBBH) (dark pink); CssS homologs with two or more cysteine residues in their extracellular domains, as predicted by DeepTMHMM (light pink). Trees were constructed using MAFFT for sequence alignment of RpoB protein sequences and FASTTREE for maximum likelihood inference and visualised in iTOL.

Next, we examined the conservation of the CutS and CssS SKs beyond the genus *Streptomyces*. Reciprocal BLAST searches in 2,936 bacterial reference genomes revealed that 39% encoded a CutS homologue and 27% encoded a CssS homologue. Of the 2,936 genomes examined, 53% encode at least one of these two SKs and 12% encode both CutS and CssS. However, further inspection of the amino acid sequences of these homologues revealed that only 6.9% of CutS and 4.8% of CssS homologues contain two or more cysteines in their extracellular domains. Phylogenetic analysis using a maximum likelihood tree based on RpoB revealed a broad distribution of these homologues across bacterial taxa (Figure 7B). The presence of both CutS and CutR with two extracellular cysteines was universally conserved across all tested *Streptomyces* strains, with only three additional genomes, all from actinobacteria, also containing both. This may reflect the fact that actinobacteria are more likely to use disulfide bond formation as a mechanism to fold their secreted proteins (32). Taken together, our findings suggest that extracellular redox sensing via conserved cysteine residues in the sensor domains of sensor kinases (SKs) may represent a broadly important mechanism across the bacterial kingdom. While CutS and CssS appear to be largely restricted to the genus *Streptomyces*, this sensing function is likely carried out by a diverse array of SKs in other bacterial lineages.

## DISCUSSION

In this study, we demonstrate that loss of the highly conserved CutRS TCS in *Streptomyces venezuelae* leads to dramatic changes in both growth and antibiotic production. To investigate the underlying mechanisms, we first defined the *S. venezuelae* CutR regulon. This revealed two conserved CutR targets—*htrA3* and *htrB*—shared between *S. venezuelae* and the distantly related *S. coelicolor*. These genes encode members of the high temperature requirement A (HtrA) family of proteases. In *S. venezuelae*, CutR represses *htrB* and activates *htrA3*. CutR also activates *vnz_08815*, a gene predicted to encode a cell wall amidase of unknown function. Using ReDCaT-SPR, we identified a consensus CutR binding motif (TAWATAAA) in the promoters of *htrA3* and *vnz_08815*, which was also found in five additional CutR-enriched promoter regions, thereby defining the complete CutR regulon. We further showed that the position of CutR binding relative to transcription start sites (TSSs) correlates with transcriptional outcome—either activation or repression. TMT proteomic analysis confirmed these regulatory effects, showing an increase in the levels of HtrB and a decrease in HtrA3 levels in the absence of *cutRS*. Given that CutRS coordinates the expression of two of the four conserved *htrA*-like genes in *S. venezuelae* and *S. coelicolor*, we conclude that CutRS functions in the secretion stress response, likely acting in parallel with the CssRS TCS.

While the CssR regulon remains undefined, previous work shows that CssR activates *htrA1*, *htrA2*, and *htrB* while having no effect on *htrA3* (33). Thus, CutRS and CssRS appear to act in opposition: they regulate distinct subsets of *htrA*-family genes and exert opposing effects on *htrB* expression. HtrA proteins support extracellular protein folding and degrade irreparably misfolded proteins in both Gram-negative and Gram-positive bacteria (34). CssRS, conserved in Gram-positive species, is homologous to the envelope stress-sensing system CpxAR in *E. coli*. Our bioinformatic analysis shows that while CutRS and CssRS are widely conserved in *Streptomyces*, most other Actinobacteria encode one or the other, and homologs are also present across other bacterial phyla (Figure 7).

Proper folding of secreted proteins requires disulfide bond formation, catalyzed by DsbA-family proteins located on the extracellular face of the cytoplasmic membrane (35). The discovery that the sensor domain of CutS contains two conserved cysteines suggests a redox-based sensing mechanism. Indeed, mutating these residues to serines (C85S, C103S) led to constitutive activation of CutR, with elevated *htrA3* expression and enhanced repression of *htrB*. This supports a model in which CutS monitors disulfide bond formation as a proxy for protein folding quality in the extracytoplasmic space.

Intriguingly, the extracellular domain of CssS also contains two conserved cysteines, and CutS and CssS are the only conserved *Streptomyces* sensor kinases (SKs) with this feature. This suggests that CssS may similarly sense disulfide bond status, which we intend to explore in future work. Moreover, *cssRS* is overexpressed in the Δ*cutRS* mutant, and deleting *cssRS* in this background restores wild-type growth, further supporting antagonistic functions for these two systems. Additional work is needed to define the CssR regulon and elucidate how CutRS and CssRS jointly regulate the HtrA proteases.

To our knowledge, this is the first detailed proposal of a mechanistic sensing model for a bacterial secretion or envelope stress response system. *Streptomyces* species are common in soil and the plant rhizosphere and endosphere (36), where protein secretion plays a vital role in breaking down complex organic material for nutrient acquisition. These environments are also oxygen-variable. We propose that, as illustrated in Figure 8, CutS senses oxygen availability via disulfide bond formation. Under anaerobic or microaerobic conditions, disulfide bonds cannot form, leading to an accumulation of misfolded extracellular proteins. We suggest that a disulfide bond fails to form in CutS under these conditions, activating the CutRS system and promoting expression of *htrA3*, which aids in resolving misfolded proteins.

**Figure 8.**
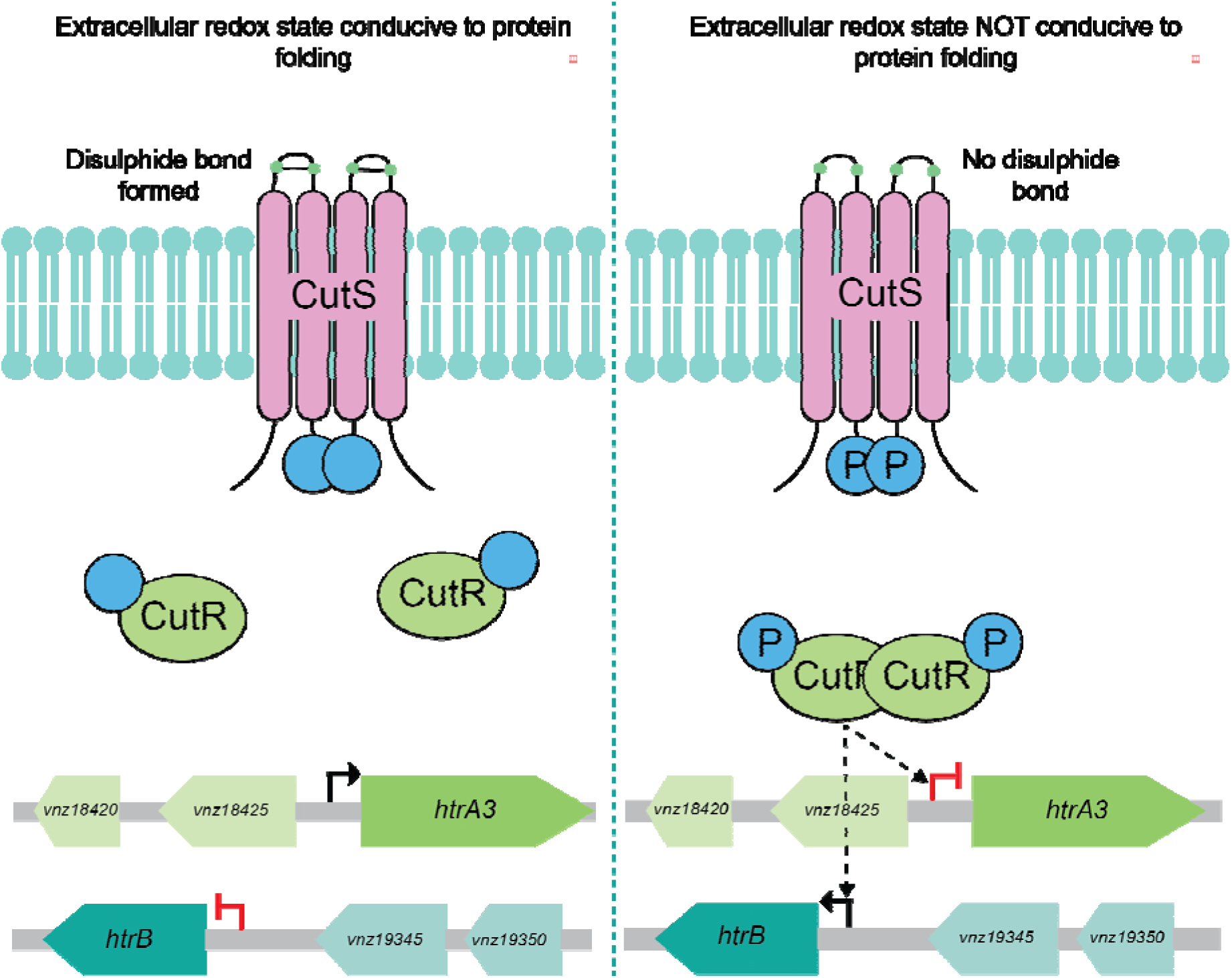
| A proposed model for the extracellular redox-sensing activity of the CutRS system in Streptomyces spp. When the extracellular space is conducive to protein folding (left side) the cysteine residues in the extracellular sensor domain of the SK CutS (pink) form a disulphide bond. In this state the CutRS system is inactive, htrA3 is expressed and htrB is repressed. When the extracellular space in not conducive to protein folding (right side) the disulphide bond in the extracellular sensor domain of CutS is unable to form. In this state the SK CutS is activated, autophosphorylates and transfers this phosphate to the cognate RR CutR (light green). Phosphorylated CutR homodimerizes and binds DNA, repressing the transcription of htrA3 and activating htrB. These genes encode dual function chaperone / proteases which can act to restore the cell to homeostasis.

Oxygen sensing via disulfide bond status has precedence. In *Shewanella* species, the MtrABC porin-cytochrome complex facilitates extracellular electron transfer under anoxic conditions (37). The periplasmic MtrC subunit contains a conserved CX₈C motif that forms a redox-active disulfide only in the presence of oxygen, allowing the cell to modulate reactive oxygen species and electron flux (38). Similarly, in *Pseudomonas aeruginosa*, the periplasmic protein YfiR contains four redox-active cysteines, two of which form a disulfide under oxidative conditions. YfiR modulates the activity of the diguanylate cyclase YfiN in response to oxygen availability (39).

The widespread conservation of sensor kinases with extracellular cysteines across diverse bacterial phyla suggests that redox and oxygen sensing outside the cell is a fundamental feature of bacterial stress response systems. Future research into these pathways may reveal new regulatory strategies for secretion stress, redox adaptation, and virulence in a broad range of bacterial species.

## METHODS

### Growth media and supplements

Protocols for culturing and genetically manipulating *Streptomyces* species are all freely available from http://actinobase.org (40). Strains were grown in the following media (per L volume): Yeast Peptone (YP) agar (10 g yeast extract, 20 g bacto-peptone, 20 g agar); YPD (YP + D-glucose) agar (YP agar with the addition of 100 ml of a 40% filter-sterilized D-glucose); 2xYT (16 g tryptone, 10 g yeast extract, 5 g NaCl); SFM agar (20 g soy flour, 20 g mannitol, 20 g agar); lysogeny broth (LB) (10 g tryptone, 5 g yeast extract, 10 g NaCl) and LB soft agar (LB with the addition of 20 g agar). When appropriate, the media was supplemented with antibiotics at the following concentrations: hygromycin (50 µg/ml), apramycin (50 µg/ml) and nalidixic acid (25 µg/ml). NaCl was excluded from LB when hygromycin was used for selection.

### Strains, plasmids, and primers

The bacterial strains, plasmids and primers used or generated in this study are listed in Supplementary Tables 2, 3 and 4, respectively.

### Strain generation

Deletion mutants were complemented using the native gene and promoter cloned into the phage integrative vector pSS170 (40) using Gibson Assembly. Overexpression strains were generated by cloning the gene(s) of interest into pIJ10257 ^28^ using Gibson Assembly. CRISPR mediated gene deletions were carried out using pCRISPomyces2 as per the protocol described by Cobb et al.(41). In all cases the resulting construct was then used to transform *E. coli* ET12567/pUZ8002 (42, 43) and conjugated into the mutant strains as follows. Single colonies of ET12567/pUZ8002 containing the complementation construct were inoculated into 10 ml LB with appropriate selection and incubated overnight at 37°C with shaking at 220 rpm. The overnight culture was sub-cultured 1:100 in 50 ml LB with appropriate selection and grown to OD_600_ ∼0.6. The cultures were washed with 10 ml ice-cold LB twice to remove the antibiotic and finally resuspended in 1 ml ice-cold LB. 500 µl of the *E. coli* cell suspension was mixed with 20 µl of *S. venezuelae* spores by inversion and then pelleted by centrifuging at 15,871 xg for 2 minutes. The supernatant was removed, and the pellet resuspended in 150 µl residual supernatant. Serial 10-fold dilutions were plated on SFM and incubated at room temperature for 16-20 hours, overlaid with 1 ml sterile distilled H_2_O containing appropriate selection and then incubated at 30°C for 3-7 days until colonies appeared. Colonies were re-streaked on SFM agar with appropriate selection at least once before being plated for spore preparation.

### Generation of S. venezuelae strain M1702

Deletion of the chloramphenicol biosynthetic gene cluster (BGC) and jadomycin BGC was achieved using the meganuclease system (44). The upstream flank for the chloramphenicol BGC was amplified with oligonucleotides NH112ChlPCR1F and NH115ChlPCR2R, whereas the upstream flanking of the jadomycin cluster was amplified with NH120JadPCR1F and NH123JadPCR2R. The upstream flanking sequences were then cloned into the BgII and HindIII sites of pIJ12738 (44). The downstream flanking sequence of the chloramphenicol BGC was amplified with oligonucleotides NH116ChlPCR3F and NH119ChlPCR4R, whereas the upstream flanking sequence of the jadomycin BGC was amplified with NH124JadPCR3F and NH127JadPCR4R. The downstream flanking sequences were combined with the corresponding upstream flanking sequences in pIJ12738 by cloning into the HindIII and SpeI sites. These pIJ12738 derivatives containing the flanking sequencing to each cluster followed by a I-SceI meganuclease cut site were then mobilized via conjugation through *E. coli* ET12567/pUZ8002 and single crossover recombinants were selected for with apramycin. We then conjugated pIJ12742 containing the meganuclease into the single crossover recombinants. Expression of the meganuclease results in a double strand break, which can be resolved by a second crossover event. Subsequent double crossovers were screened by PCR to determine if they were gene cluster deletion mutants or if they had reverted to wild-type. Gene cluster deletion mutants were incubated at 37°C for two generations to promote loss of pIJ12742. The resulting strains were Δ*cml* (M1700) and Δ*jad* (M1701). M1700 was used as the starting material to introduce the jadomycin BGC deletion derivative of pIJ12738, and single crossover recombinants were used to introduce the meganuclease (on pIJ12742) followed by screening for loss of the jadomycin BGC and removal of the plasmid, generating the final double mutant of Δ*cml*Δ*jad* (M1702).

### Antimicrobial bioassays

5 µl of spores of the relevant *S. venezuelae* strains were inoculated onto the centre of a YP or YPD agar plate and incubated for 5 days at 30°C. The *B. subtilis* indicator strain was grown in 10 ml LB at 37°C with 220 rpm shaking the night prior to the assay. Following overnight incubation, the indicator strains were sub-cultured 1:100 into 50 ml LB and grown to OD_600_ ∼0.6. and then diluted 1:10 into 50°C Soft LB agar. 5 ml of this inoculated soft agar was then pipetted over the *S. venezuelae-*containing plate ensuring complete coverage. The plates were air-dried and subsequently incubated at 37°C overnight before imaging.

### Biomass and glucose quantification assays

5 µl of spores of the relevant *S. venezuelae* strains were inoculated on cellophane-covered YP or YPD agar plates for 5 days at 30°C. Following incubation, the cellophane disks were removed, and the mycelial mass was measured using an analytical balance. For glucose extraction the underlying agar was chopped into small pieces using a sterile scalpel and transferred to a 250 ml conical flask. 20 ml sterile distilled H_2_O was added, and samples were incubated for 2 hours at 25°C with shaking at 220 rpm. The resulting liquid was aspirated from the flask and diluted 1:1000 with sterile distilled H_2_O before being assayed using the Glucose (GO) Assay Kit (Sigma Aldrich) following manufacturer’s instructions.

### Chromatin immunoprecipitation followed by sequencing (ChIP-Seq)

5 µl of spores of the relevant *S. venezuelae* strains were inoculated onto cellophane-covered YP or YPD agar plates in triplicate and grown for 5 days at 30°C. To cross-link proteins to DNA the cellophane disks were removed and submersed in 10 ml fresh 1% (v/v) formaldehyde solution at room temperature for 20 minutes. Following a 5-minute incubation in 10 ml 0.5 M glycine solution the mycelium was harvested and washed twice with 25 ml ice-cold PBS (pH 7.4) before being flash frozen in liquid nitrogen and stored at – 80°C. For lysis, the pellets were resuspended in 2 ml lysis buffer (10 mM Tris-HCl pH 8.0, 50 mM NaCl, 10 mg/ml lysozyme, EDTA-free protease inhibitor) before 750 µl was transferred to a 2 ml microcentrifuge and samples incubated at 37°C for 30 minutes. For fragmentation 750 µl IP buffer (100 mM Tris-HCl pH 8.0, 250 mM NaCl, 0.5% v/v Triton X-100, 0.1% SDS, EDTA-free protease inhibitor) was added and samples sonicated on ice at 50 Hz for 20 cycles of 10 seconds with at least 1 minute of rest between each cycle. 25 µl of the crude lysate was combined with 75 µl TE Buffer (10 mM Tris-HCl pH 8.0, 1 mM EDTA) and extracted with 100 µl phenol:chloroform. 2 µl of 1 mg/ml RNase A was combined with 25 µl of the resulting extract, incubated at 37°C for 30 minutes and run on a 1% agarose gel to confirm a smear centred at 500 bp. The remaining crude lysate was centrifuged for 15 minutes at 15,871 xg at 4°C and supernatant saved. 750 µl of α-FLAG M2 magnetic beads (Sigma Aldrich) were washed in 3.75 ml 0.5% IP buffer (IP buffer – 150 mM NaCl, 1% Triton X-100, 50 mM Tris-HCl (pH 8.0), 2 mM EDTA, and Roche cOmplete^™^, Mini, EDTA-free Protease Inhibitor Cocktail tablet(s)) to prepare for them binding. The bead slurry was then mixed with 40 µl clarified lysate and incubated overnight on a vertical rotator at 4°C. The lysate was removed, and the beads washed four times with 500 µl 0.5 IP buffer with 10 minutes vertical rotation at 4°C each cycle. Washed beads were incubated in 100 µl elution buffer (50 mM Tris-HCl pH 7.6, 10 mM EDTA, 1% SDS) overnight at 65°C. Eluant was removed and saved before an additional 50 µl elution buffer was added and incubated at 65°C for a further 5 minutes. Once removed 3 µl of 10 mg/ml proteinase K was added to the combined eluants and incubated at 55°C for 2 hours. DNA was extracted with 200 µl phenol:chloroform and purified using a QIAquick PCR Purification Kit (QIAGEN) following the manufacturer’s instructions. DNA was eluted in 50 µl Buffer EB (10 mM Tris-HCl pH 8.5) and 47 µl snap frozen in liquid nitrogen and stored at –80°C. The remaining 3 µl of sample was used to determine the DNA concentration and quality. The Qubit Fluorometer 2.0 with the high-sensitivity kit was used for DNA concentration whilst DNA purity was quantified using a Nanodrop™ 2000 UV-Vis Spectrophotometer. The stored DNA samples were the shipped to Novogene on dry ice for sequencing using the NovaSeq 6000 Sequencing System (Illumina). Raw sequencing data were received as FASTQ files from Novogene and processed as described previously (27). Raw files were aligned to the *S. venezuelae* NRRL B-65442 genome and enrichment normalised to a moving window of 30 nucleotides compared to a region of the surrounding 3000 nucleotides with the window moving in steps of 15 nucleotides.

### Protein overexpression and purification

Protein preparation for ReDCaT SPR was performed as follows. N-terminally 6xHis-tagged CutR was expressed from the plasmid pET28a in *E. coli* Rosetta (BL21 DE3) pLysS competent cells. Overnight culture (40 ml) was used to inoculate 4 L of LB with selective antibiotics. Cultures were incubated at 37°C with shaking at 220 rpm until OD_600 ∼_0.6 before isopropyl-β-D-1-thiogalactopyranoside (IPTG) was added to a final concentration of 1lmM. Cultures were incubated for a further 4 hours then the cells were pelleted by centrifugation and stored at –80°C. Cell pellets were defrosted on ice for 30 minutes before being resuspended in 25 ml lysis buffer (20 mM Tris-HCl pH 8.0, 75 mM NaCl, 0.1% Triton X-100, 10 mg/ml lysozyme, EDTA-free protease inhibitor) and incubated at room temperature for a further 30 minutes. Suspensions were sonicated on ice 8 times at 50 Hz for 30 seconds per cycle with 1 minute off between each sonication cycle. Cell debris was removed by centrifugation at 20,600 xg for 30 minutes at 4°C and the supernatant filtered through a sterile 0.22 µm filter (Sartorius). Clarified lysate was loaded onto a 1 ml HisTrap HP column (Cytiva) pre-equilibrated with Buffer A (50 mM Tris-HCl pH 8.0, 200 mM NaCl, 5% glycerol). The loaded column was washed with increasing concentrations of imidazole (step 1: 10 mM; step 2: 20 mM; step 3: 30 mM) before the protein was eluted using 500 mM imidazole, all in the same buffer. Desired protein fractions were pooled and loaded onto a preparative-grade HiLoad 16/600 Superdex 200 pg gel filtration column (GE Healthcare) pre-equilibrated with gel filtration buffer (50 mM Tris-HCl pH 8.0, 200 mM NaCl, 10% glycerol, filtered). Desired fractions were identified and analysed for purity via SDS-PAGE before being pooled. Aliquots were flash frozen in liquid nitrogen and stored at –80°C.

### Reusable DNA Capture Technology Surface Plasmon Resonance (ReDCaT SPR)

Protein-DNA binding site scanning and footprinting was performed using ReDCaT SPR as previously described (45). Preliminary scanning was performed using promoter regions of interest divided into a series of overlapping double stranded oligonucleotide probes (Supplementary Table 4). The total length of each fragment was 40 bp including 15 bp overlaps. To create the double-stranded probes each fragment was reverse-complimented and a ReDCaT biotinylated single-stranded linker added to the 3’ end as an overhang. For each fragment DNA oligos (IDT) were resuspended to 100 µM in sterile distilled H_2_O and 55 µl forward oligo was mixed with 45 µl reverse oligo. Oligos were annealed at 95°C for 5 minutes followed by ramping to 4°C at 0.1°C/s. The annealed DNA fragments were then diluted to a final concentration of 1 µM in HBS-EP+ buffer (150 mM NaCl, 3 mM EDTA, 0.05% v/v surfactant P20, 10 mM HEPES pH 7.4). Positive hits from site scanning were selected for footprinting. 2 bp truncations were made from the 3’ end of the selected fragments and similarly from the 5’ of the inverted fragment. All ReDCaT SPR experiments were performed using a single Sensor Chip SA (GE Healthcare) on a Biacore 8K SPR system (Cytiva) as follows. For each channel FC_1_ was designated as the reference (FC_REF_) and FC_2_ as the test cell (FC_TEST_). DNA capture at 400 RUs was achieved in FC_TEST_ with a contact time of 60 seconds at a flow rate of 10 µl/min. Protein analyte (Purified CutR; 2 µM or 1 µM) was then added over both FCs for 60 seconds at a flow rate of 50 µl/min followed by a buffer only dissociation of 60 seconds. Regeneration of the chip was achieved using Regeneration buffer (1 M NaCl, 50 mM NaOH) injected for 60 seconds at a flow rate of 10 µl/min over both FCs. All experiments were run in duplicate and data analysis was performed using the Biacore Insight Evaluation software (Cytiva) and Microsoft Excel.

### Tandem Mass Tagging proteomic analysis

5 µl of spores of the relevant *S. venezuelae* strains were inoculated on cellophane-covered agar plates as triplicate spots in duplicate. After 2-or 9-day incubation at 30°C the mycelium was scraped off the cellophanes into a 15 ml falcon and resuspended in 10 ml cell lysis buffer (50 mM TEAB buffer pH 8.0, 150mM NaCl, 2% SDS, EDTA-free protease inhibitor, PhosSTOP phosphatase inhibitor). The suspension was disrupted via French press three times before being boiled at 100°C for 10 minutes. Samples were sonicated at 50 Hz four times for 20 seconds per cycle and then pelleted at 3,220 xg for 30 minutes. Protein concentration was determined using the BCA assay (Thermo Fischer Scientific) and 1 mg of protein from each sample transferred to a fresh 15 ml falcon tube. The proteins were precipitated from the supernatant with chloroform-methanol (46) and washed with acetone.

The air-dried pellets were dissolved in 50-100 µl of 2.5% sodium deoxycholate (SDC; Merck). After quantification by BCA assay, 100 µg of protein was reduced, alkylated, and digested with trypsin in the presence of 0.2 M EPPS buffer (Merck) and 2.5% SDC according to standard procedures. After digestion, the SDC was precipitated by adjusting to 0.2% TFA, and the clarified supernatant subjected to C18 solid phase extraction (SPE; OMIX tips; Agilent). Tandem mass tagging (TMT) labelling was performed using a Thermo TMT10plex™ kit (lot TL274393) according to the manufacturer’s instructions with slight modifications; the dried peptides were dissolved in 90 µl of 0.2 M EPPS buffer/10% acetonitrile, and 200 µg TMT10plex reagent dissolved in 22 µl of acetonitrile was added. Samples were assigned to the TMT channels in an order avoiding channel leakage between different samples, if possible, as detailed by Brenes et. al (47). After labelling, aliquots of 1.7 µl from each sample were combined and analysed on the mass spectrometer (detailed below) to check labelling efficiency and estimate total sample abundances. The main sample aliquots were then combined correspondingly and desalted using a 50 mg C18 Sep-pak cartridge (Waters). The eluted peptides were dissolved in 500 µl of 25 mM NH_4_HCO_3_ and fractionated by high pH reversed phase HPLC. Using an ACQUITY Arc Bio System (Waters), the samples were loaded to a Kinetex® 5 µm EVO C18 100 Å LC Column 250 x 4.6 mm (Phenomenex). Fractionation was performed with the following gradient of solvents A (water), B (acetonitrile), and C (25 mM NH_4_HCO_3_) at a flow rate of 1 ml/min: solvent C was kept at 10% throughout the gradient; solvent B: 0-5 min: 5%, 5-10 min: 5-13% curve 5, 10-70 min: 13-40%, 70-75 min: 40-50%, 75-80%: 50-80%; followed by 5 min at 80% B and re-equilibration to 5% for 24 minutes. Fractions of 1 ml were collected, dried down, and concatenated to produce 20 final fractions for MS analysis. Aliquots of 30-40 µl were removed and stored at –80°C for global proteome quantification. Aliquots of all concatenated fractions were analysed by nanoLC-MS/MS on an Orbitrap Fusion™ Tribrid™ mass spectrometer coupled to an UltiMate® 3000 RSLCnano LC system (Thermo Fisher Scientific). The samples were loaded onto a trap column (nanoEase M/Z Symmetry C18 Trap Column, Waters) with 0.1% TFA at 15 µl/min for 3 minutes. The trap column was then switched in-line with the analytical column (nanoEase M/Z column, HSS C18 T3, 1.8 µm, 100 Å, 250 mm x 0.75 µm, Waters) for separation using the following gradient of solvents A (water, 0.1% formic acid) and B (80% acetonitrile, 0.1% formic acid) at a flow rate of 0.25 µl/min; for global protein analysis: 0-4 min 3% B (parallel to trapping); 4-17 min increase B curve 4 to 14%; 17-80 min linear increase B to 35%; 80-102 min linear increase B to 55% followed by a ramp to 99% B and re-equilibration to 3% B for 23 minutes; for enriched phosphopeptides: 0-4 min 3% B (parallel to trapping); 4-11 min increase B curve 4 to 12%; 11-52 min linear increase B to 43%; 52-61 min linear increase B to 55%, 61-64 min linear increase B to 65%, followed by a ramp to 99% B and re-equilibration to 3% B for 23 minutes. Data were acquired with the following parameters in positive ion mode: MS_1_/OT: resolution 120K, profile mode, mass range m/z 400-1800, AGC target 2e5, max inject time 50 ms; MS_2_/IT: data dependent analysis with the following parameters: top10 in IT Turbo mode, centroid mode, quadrupole isolation window 1.6 Da, charge states 2-5, threshold 1.9e4, CE=35, AGC target 2e4, max. inject time 70 ms, dynamic exclusion 1 count/7 s with ±7 ppm, multistage activation on with neutral loss 97.98 Da (for phosphopeptides only); MS_3_ synchronous precursor selection (SPS): 10 SPS precursors, isolation window 1.6 Da, HCD fragmentation with CE=65, Orbitrap resolution 50k, AGC target 1.5e5, max inject time 175 ms.

The acquired raw data were processed and quantified in Proteome Discoverer 3.1 (Thermo Fisher Scientific); all mentioned tools of the following workflow are nodes of the proprietary Proteome Discoverer (PD) software. The *Streptomyces venezuelae* (http://strepdb.streptomyces.org.uk/, 7420 entries) protein fasta database was imported into PD adding a reversed sequence database for decoy searches; a database for common contaminants (maxquant.org, 245 entries) was also included. For global protein quantification, the database search was performed using the incorporated search engine CHIMERYS (MSAID, Munich, Germany). The processing workflow also included spectrum selection, the TopN Peak Filter with 20 peaks per 100 Da, and the reporter ion quantification by most confident centroid (20 ppm). The inferys_3.0.0. fragmentation prediction model was used with fragment tolerance of 0.5 Da, enzyme trypsin with 1 missed cleavage, variable modification oxidation (M), fixed modifications carbamidomethyl (C) and TMT10plex on N-terminus and K. Identifications were calculated for False Discovery Rate (FDR) 0.01 (strict) and 0.05 (relaxed). The results were exported into a Microsoft Excel table including data for normalized and un-normalized abundances, ratios for the specified conditions, the corresponding p-values and adjusted p-values, number of unique peptides, q-values, PEP-values, identification scores from both search engines; FDR confidence filtered for high confidence (strict FDR 0.01) only. Further filtering included removal of contaminants and single unique peptide matches.

### Chemical extraction of chloramphenicol from solid media

Two 5 µl aliquots of spore stock were used to inoculate cellophane-covered 30 ml agar plates with *Streptomyces* spp. before incubation at 30°C for 2 days. Subsequently the cellophane was removed. Agar was cut into 1 cm^3^ pieces using a sterile razor blade and transferred to a 100 ml Duran bottle. 30 ml of HPLC-grade ethyl acetate was added to the agar, briefly shaken, and left at room temperature for 1 hour. The top 25 ml of the ethyl acetate was then transferred to a sterile 50 ml falcon tube and the sample was dried under vacuum. The sample was then resuspended in 500 µl HPLC-grade methanol and transferred to a 1.5 ml microcentrifuge tube. Insoluble particulates were removed via centrifugation at 15,871 xg for 10 minutes with the uppermost 200 µl of solution transferred to a 2 ml HPLC vial containing a 200 µl glass insert for downstream chloramphenicol quantification.

### Chloramphenicol Quantification by High-performance Liquid Chromatography

Analytical HPLC was performed on a 1290 Infinity II LC System (Agilent). A chloramphenicol standard curve was prepared in triplicate with serial dilutions of chloramphenicol in methanol from 1000 – 0.01 µg/ml. Chloramphenicol standards and extracted samples were chromatographed over a Kinetex 2.6 µm XB-C18 110 Å, 100 x 4.6mm column (Phenomenex) eluting with a linear gradient: mobile phase A, 0.1% (v/v) trifluoroacetic acid; mobile phase B, acetonitrile; 0 min, 10%B; 1 min, 10%B; 11 min, 100%B; 13 min, 100%B; 13.2 min, 10%B; 15 min, 10%B; flow rate 1 ml/min; injection volume 10 µL. UV absorbance was measured at 278 nm and the peak area at this wavelength was measured for analysis. The minimum concentration of chloramphenicol that could accurately be detected under these conditions was 0.5 µg/ml.

### Quantitative reverse transcription PCR (qRT-PCR)

To perform quantitative RT-PCR (qRT-PCR), *S. venezuelae* wild-type, Δ*cutRS,* Δ*htrA3* and Δ*htrA3* + pIJ10257-*htrB* strains were grown on top of sterilized cellophane discs on YP or YPD agar at 30°C for 8-10 days. Colonies were removed from the cellophanes kept in liquid nitrogen whilst being crushed using a sterile pestle and mortar on dry ice. Crushed samples were resuspended in 1 ml RLT Buffer (Qiagen) supplemented with 1 % 2-mercaptoethanol. This suspension was added to a QIA-shredder column (Qiagen) with the flowthrough transferred to a new 1.5 ml microfuge tube (leaving the pellet). Then, 700 µl acidic phenol:chloroform was added and the sample incubated at room temperature for 3 minutes and subsequently centrifuged at 15,871 xg. for 20 min. The resulting upper phase was mixed with 0.5 volumes 95l% ethanol. This was then applied to a RNeasy Mini spin column (Qiagen) and processed following the manufacturer’s protocol. The eluant was treated with the Turbo-DNase kit (Invitrogen) followed by a RNeasy mini clean-up kit (Qiagen) both used according to manufacturer’s instructions. The samples were then flash frozen in liquid nitrogen for storage at −80°C. RNA was quantified with Nanodrop and Qubit Fluorometer and∼1 mg converted to cDNA using the LunaScript RT SuperMix Kit (NEB) as per the manufacturer’s instructions, including No-RT control reactions. Primers for each transcript were optimized using serial dilutions of template DNA and checked for efficiency by generating standard curves and calculating as follows: *E=(10slope−1)–1*. Primer sets with an efficiency between 90 and 110l% and within 5 % of each other were selected. qRT-PCR was run using the QuantStudio1 (Applied Biosystems) with Luna Universal qPCR Master Mix according to the manufacturer’s guidelines for a 20 µl reaction with 2 µl template cDNA and a final concentration of 0.25 mM of each primer. Controls with no reverse transcriptase were run to ensure the absence of contaminating gDNA. ΔC_T_ values were normalized to the *hrdB* gene, which encodes the housekeeping RNA polymerase sigma factor.

### Phylogenetic analysis

Genomes for use in phylogenetic analysis were obtained from the NCBI database(48), in all cases only complete and ‘reference’ RefSeq genomes were selected to avoid duplication. The accession numbers for genomes used to generate Figure 6A and 6B are given in Supplementary Information 1 and 2 respectively. Protein sequence alignments were carried out using MAFFT version 7 (mafft –-memsavetree –-retree 1 –treeout) (49, 50). Aligned sequences were used to produce a maximum likelihood tree using FASTTREE (FastTree –jtt alignment_file > output_tree) (51). The resulting trees were annotated in Interactive Tree Of Life (iTOL) v7 (52). Homologues of SvCutS and SvCssS were identified by reciprocal BLAST search, where reciprocal BLAST best hits were used in further analysis.

### Protein Sequence Alignments

Protein sequences for alignment were obtained via reciprocal blast searches using *S. venezuelae* GCF_001886595.1 CutS and CssS as the query sequences. The sequences obtained for CutS and CssS homologues are available in Supplemental information 1.7 and 1.8 respectively. Sequences were aligned with Clustal Omega (53) and visualised in Jalview (54).

### Data availability

The ChIP-seq data generated in this study have been deposited in the GEO database under the accession code GSE225370. The proteomics data generated in this study have been deposited in the ProteomeXchange database with identifier PXD051851. The genome sequence accession numbers and protein identifiers used to generate the data in Figure 7A and 7B available as Supplementary Information 1 and 2 respectively. Protein identifiers used to generate Supplementary Figure 2 and Supplementary Figure 4 are available as Supplementary Information.

## Acknowledgements

We would like to thank Professor Mervyn Bibb (John Innes Centre) for his contributions to the conceptualisation and generation of *S. venezuelae* strain M1702. We also thank past and present members of the Hutchings and Wilkinson groups (John Innes Centre) and Dr Alison MacFadyen (The Sainsbury Laboratory) and Professor Tracy Palmer (Newcastle University) for useful discussions. We thank John Innes Centre support services for growth media, glassware, plasticware and waste disposal and Phil Robinson for his photography expertise. This work was supported by the BBSRC doctoral training programme grant BB/M011216/1 (T.C.M.); by the BBSRC-funded Institute Strategic Programme Project BBS/E/J/000PR9790 to the John Innes Centre and by the BBSRC-funded Institute Strategic Program Harnessing Biosynthesis for Sustainable Food and Health (HBio) grant BB/X01097X/1 to the John Innes 900 Centre.

## Author Contributions

Conceptualization – T.C.M. and M.I.H.

Data curation – A.D.M.B., G.C., C.M. and G.S.

Formal analysis – T.C.M., A.D.M.B., G.C., C.M. and G.S.

Funding acquisition– M.I.H. and B.W.

Investigation – T.C.M., A.D.M.B., N.A.H., J.C., G.C., C.M. and G.S.

Methodology – T.C.M., A.D.M.B., G.C., C.M. and G.S.

Project administration – T.C.M., M.I.H. and B.W.

Resources – C.M., G.S., M.I.H. and B.W.

Software – G.C., C.M. and G.S.

Supervision – M.I.H. and B.W.

Validation – T.C.M. and A.D.M.B.

Visualization– T.C.M. and A.D.M.B.

Writing – original draft – T.C.M., A.D.M.B. and M.I.H.

Writing – review and editing – T.C.M., A.D.M.B., G.C., C.M., G.S., B.W. and M.I.H.

## SUPPLEMENTARY FIGURES

**Supplementary Figure 1.**
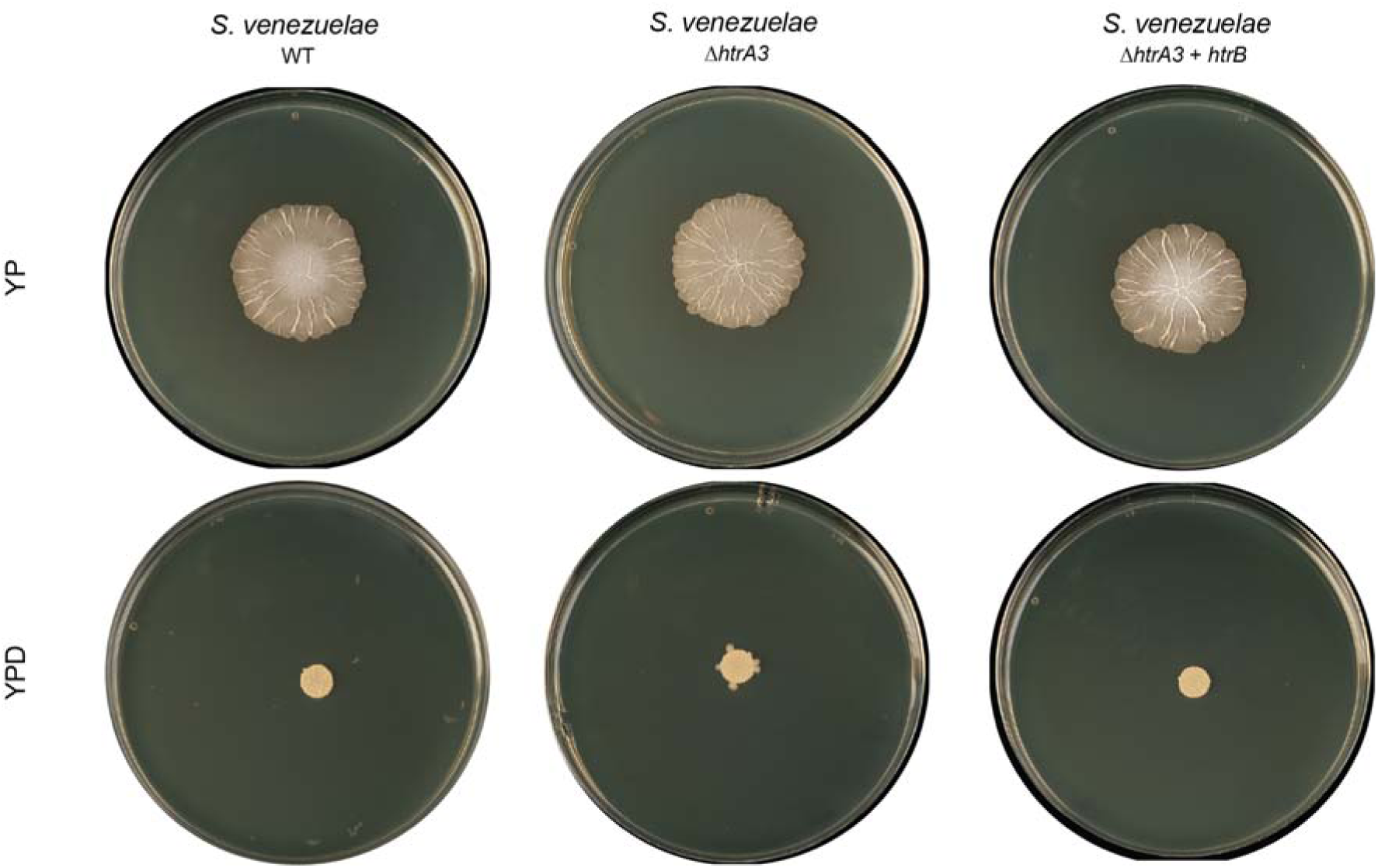
| Overexpression of htrB in the S. venezuelae ΔhtrA3 background is unable to reproduce the ΔcutRS phenotype. A. S. venezuelae wild-type, ΔhtrA3 and ΔhtrA3 with htrB overexpression colonies grown on YP (-glucose) and YPD (+ glucose) for 10 days at 30°C.

**Supplementary Figure 2.**
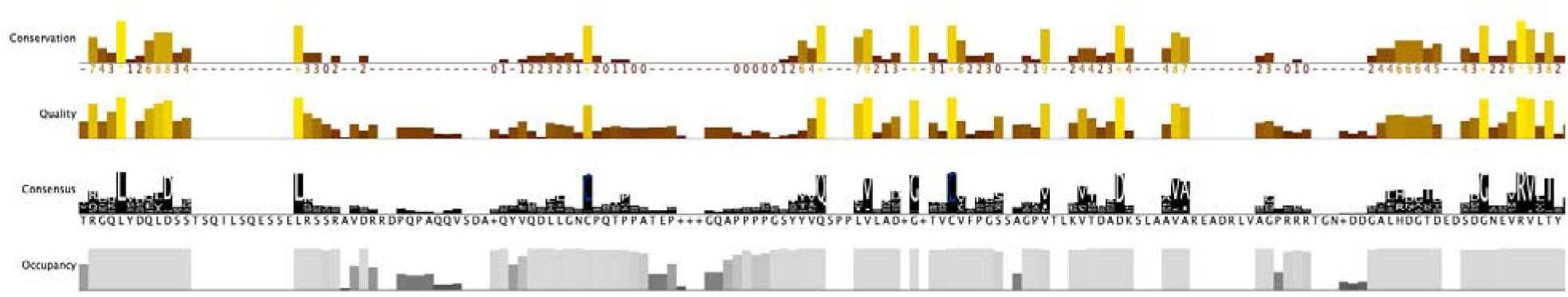
| Residue conservation in the extracellular sensor domain of CutS. Clustal OMEGA alignment of 105 CutS homologous from Streptomyces spp. showing two highly conserved cysteine residues in the extracellular sensor domain. Alignment view taken from Jalview giving the alignment conservations, quality, consensus and occupancy over one section of the alignment. Sequences use to generate this figure can be found as Supplementary File 3.

**Supplementary Figure 3.**
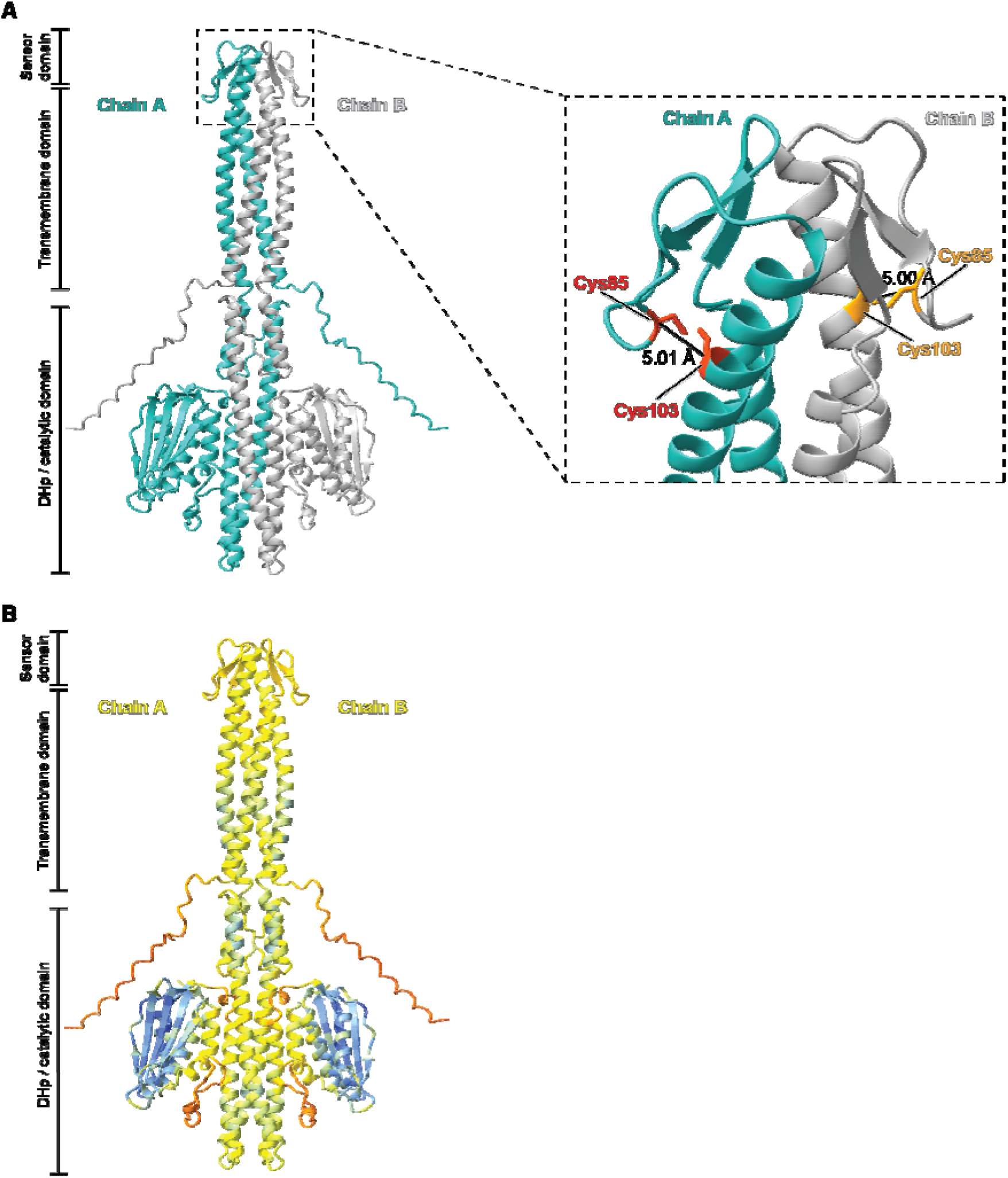
| CutS is predicted to form disulphide bonds between the two cysteine residues present in the extracellular sensor domain. A. AlphaFold2-predicted structure of homodimeric S. venezuelae CutS. The extracellular sensor domain, transmembrane domain and DHp / catalytic domain are annotated. (Inset) Close-up view of the predicted extracellular sensor domain highlighting the intra-chain disulphide bonds and predicted bond lengths. Chain A: green; Chain B: grey. B. AlphaFold2-predicted structure of homodimeric S. venezuelae CutS coloured by confidence score (pIDDT): very high (pIDDT > 90; dark blue), confident (90 > pIDDT > 70; light blue), low (70 > pIDDT > 50; yellow) and very low (pIDDT < 50; red).

**Supplementary Figure 4.**
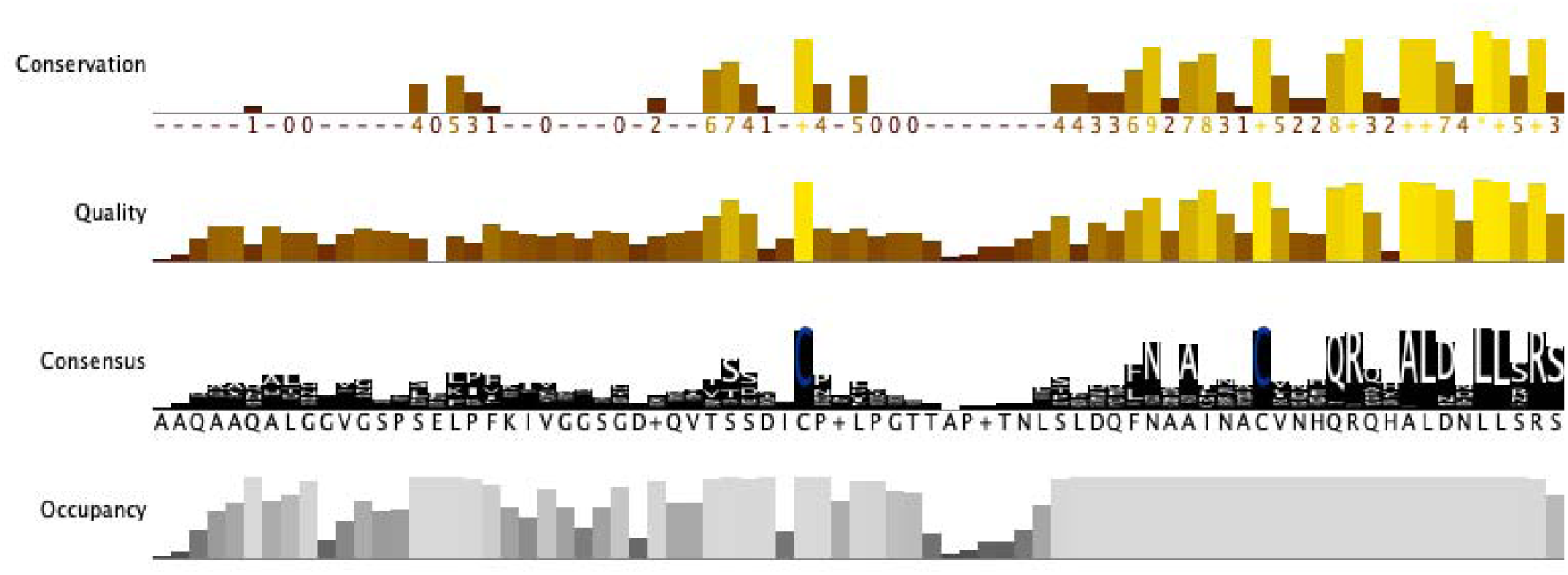
| Residue conservation in the extracellular sensor domain of CssS of Streptomyces spp. Clustal OMEGA alignment of 105 CssS homologous from Streptomyces species showing two highly conserved cysteine residues in the extracellular sensor domain. Alignment view taken from Jalview giving the alignment conservations, quality, consensus and occupancy over one section of the alignment. Sequences use to generate this Figure can be found as Supplementary File 3

